# Revisiting the erythroid-myeloid lineage decision – a data-driven dynamical model analysis

**DOI:** 10.1101/197822

**Authors:** Jose Teles, Victor Olariu, Carsten Peterson

**Author notes:** These authors contributed equally to this work (Jose Teles, Victor Olariu).

## Abstract

It is widely conjectured that the mutually antagonizing pair of transcription factors GATA1 and PU.1, determines the choice between the erythroid and myeloid lineages in hematopoiesis. In theoretical approaches, this appears natural with a bistable switch driving the decision. Recent extensive binding and gene expression experiments with some focus on the triad GATA1, GATA2 and PU.1 indicate that GATA2 may be more involved in this lineage decision than previously anticipated. Here, we analyze these experimental data by modeling regulatory sub-networks with deterministic rate equations. Using network dynamical parameters determined by the data, we deduce from increasing the self-interaction bindings *in silico* among the triad genes that GATA2 and PU.1 exhibit non-linear behavior with one unstable and one stable state. This is in contrast to GATA1, which shows smoother behavior. We extend the network to include the downstream regulators FOG1 and CEBPA, and extract the nature of the corresponding regulatory interactions, excitatory or suppressing, between this pair and the triad by fitting to experimental gene expression time series. Based on this extended network, we simulate and explore different knockout scenarios, providing insight into the role of these regulators in the process of lineage specification, as well as predictions for future experimental validation. We address the mechanism of GATA switching as a mechanism of lineage differentiation by investigating the dynamics of FOG1 regulation by GATA2 and GATA1. Overall, this analysis strongly suggests that within this network, GATA2 is the key driver of erythroid lineage specification through its repression of PU.1, whereas GATA1 appears to be more relevant for the downstream differentiation events.

## Introduction

The transcription factors (TFs) GATA1, GATA2 and PU.1 are important regulators in hematopoiesis. Their regulatory interactions and dynamic expressions provide an attractive system for understanding mechanisms of lineage specification and differentiation [1]. GATA2 is associated primarily with stem cells and multipotent progenitors, GATA1 with erythroid cells and megakaryocytes, and PU.1 with myeloid and lymphoid cells (reviewed in [2, 3]). Recently, the importance of these TFs in driving commitment and differentiation of multipotent cells has been accessed through genome-wide experiments performed on the FDCPmix cell line [4]. In this study, global gene expression profiles were generated throughout the unilineage specification and differentiation of hematopoietic multipotent cells to two opposing cell fates, erythroid and neutrophil. In addition, global ChIPSeq analysis of GATA1, GATA2 and PU.1 in multipotent and differentiated cells was also performed. The transition from multipotent to the erythroid and myeloid lineages has also been the subject of computational studies. In particular the self-activation and mutual antagonism interactions in the GATA1/PU.1 pair have been considered a paradigmatic system for lineage specification [5, 6]. The properties of the system that lead to bistability and support the existence of a primed progenitor state have been explored, both through pure theoretical efforts [7, 8] as well as approaches based on experimental data [9]. A computational analysis was also ventured in [4], expanding the GATA1/PU.1 switch to GATA2 with the aim of establishing the nature of unknown interactions between these three regulators, excitatory and repressive, provided by ChIPSeq analysis. Exploration of all 32 possible combinations of positive and negative interactions between GATA2, GATA1 and PU.1 was performed by comparing against dynamical models fitted to the time series, leading to the inference of the regulatory logic for the GATA1/GATA2/PU.1 sub-network. In this approach, the dynamic (time dependent) nature of the binding data was an important factor in obtaining the solutions. GATA2 repression of PU.1 was a consistent feature of all good solutions, suggesting that PU.1 repression by GATA2 is central to erythroid lineage specification. Beyond lineage specification events, the regulatory mechanisms by which gene expression programs coordinate throughout differentiation have also been an active topic of research. A common view is that master regulators involved in the decision switch not only regulate each other but also affect the expression of a large number of target genes downstream, activating transcriptional programs affiliated with their own lineage and repressing programs affiliated with alternative fates [10]. This mechanism has recently been described for a core transcriptional network in erythroid differentiation [11]. The complexity of cross-regulatory interactions influencing erythroid lineage specification and differentiation is also well illustrated by the process of GATA switching, where GATA2 is gradually replaced by GATA1 in binding to its target genes, relinquishing control of the transcriptional program driving differentiation [12, 13].

In this work, we address fundamental questions of early erythroid lineage specification and differentiation, by expanding the analysis in [4] along three main directions:

- With the GATA2-PU1 axis as a potential important factor for the commitment transition, we revisit the question of bifurcation structures in the earlier stages using the determined architecture and bindings. We do this by varying the self-interactions one by one for GATA1,GATA2 and PU.1 with results pointing at GATA2 being a key driver in erythroid commitment.
- Using the same method as in [4] we expand the triad sub-network into the immediate downstream components FOG1 and CEBPA, affiliated with the erythroid and myeloid lineages, respectively. We seek to determine the nature of these interactions by exploring the different combinations of unknown regulatory signs. Our results suggest that PU.1 represses FOG1 and promotes CEBPA, whereas GATA2 promotes FOG1. Taken together, these findings support the notion that mutual antagonism extends to lower levels of the network.
- Departing from this extended network composed of GATA1, GATA2, PU.1, FOG1 and CEBPA, we perform *in silico* knockout experiments, which provide insight into mechanisms of lineage specification within the core triad network, as well as into regulatory processes involved in early differentiation, such as GATA switching.

## Results

### Bifurcation properties of the triad show no bistability for GATA1

In our previous work, we have been able to infer regulatory interactions of the GATA1/GATA2/PU.1 triad, using dynamical modeling based on experimental gene expression and binding data. The reconstructed network (Figure 1A) expands to GATA2 the well-known interactions in the GATA1-PU.1 axis and suggests that repression of PU.1 by GATA2 is a key interaction in the system.

**Figure 1:**
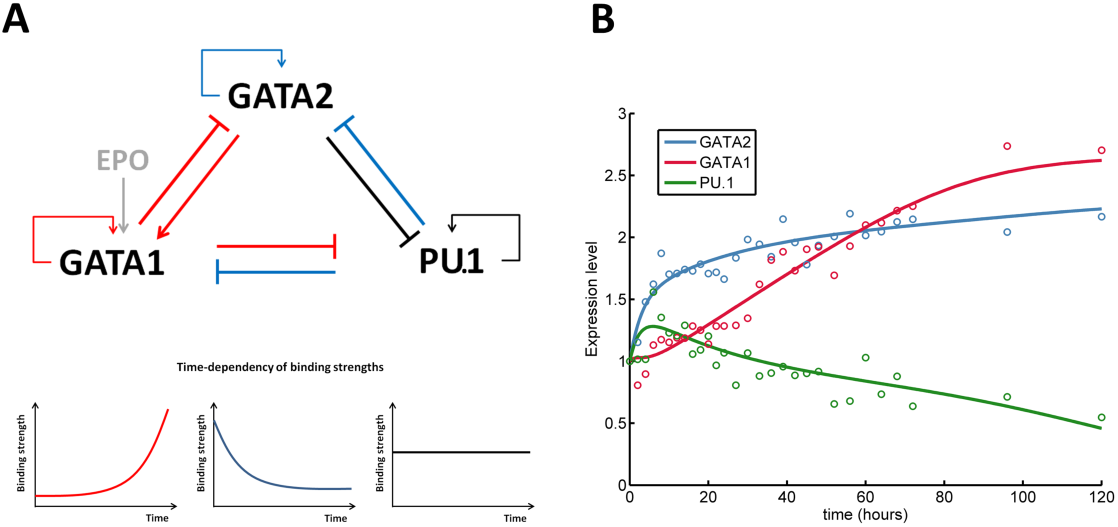
The triad interaction circuit depicting the most likely architecture from fitting to gene expression data. (A) The core network for the mutual and self-regulatory interactions between GATA1, PU.1 and GATA2. Red interaction indicates increase in time of the binding strength, blue corresponds to decreasing and black shows constant binding strengths (as shown in the panel below the network diagram). The external signal Epo is represented in grey. (B) Expression time series of GATA1 (red), PU.1 (green) and GATA2 (blue) from the network model (solid lines) with the best set of estimated parameters. Experimental data are represented by circles. Expression levels are normalized by expression at t=0 (multipotent state).

Based on this architecture, our model is able to accurately recapitulate experimental gene expression time-course data, exhibiting up-regulation of GATA1 and down-regulation of PU.1, typical of erythroid differentiation, as well as the quick up-regulation up to a plateau level for GATA2 (Figure 1B). In the current work, we further explore the impact of each of these regulators in the process of lineage specification by performing bifurcation studies and observing the qualitative changes in the stability of attractors of the dynamical system describing the molecular interactions within the network. We computed the steady state solutions as functions of the self-interaction binding strengths of each gene. When varying the self-interaction strength for GATA1, the bifurcation diagram shows no significant non-linear behaviour. All three genes display one stable attractor for all values of the varied parameter (Figure 2A).

**Figure 2:**
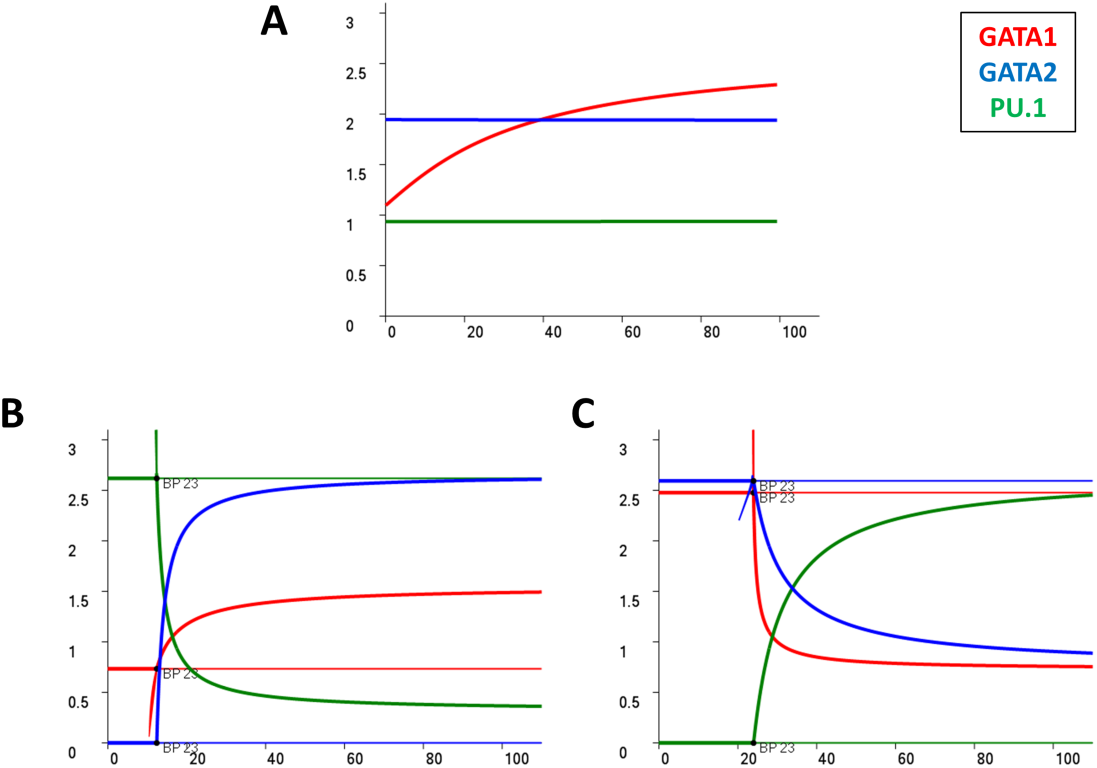
Steady state analysis of the transcription circuit by varying the self-interaction strengths of the three genes. (A) The steady state values of GATA1, PU.1 and GATA2 as a function of GATA1 self-interaction strength. All three genes exhibit one stable attractor at almost constant values. (B) The steady state values of GATA1, PU.1 and GATA2 as a function of GATA2 self-interaction strength. For low values of varying parameter, GATA1 and GATA2 have one stable attractor at low values while PU.1 has one stable attractor at high values. At the bifurcation point the GATA1 and GATA2 factors have one stable attractor at higher values and one unstable one at low values while PU.1 has an unstable attractor at high values and a stable one at low values. (C) The steady state values of GATA1, PU.1 and GATA2 as a function of PU.1 self-interaction strength. For low values the PU.1 self-interaction, GATA1 and GATA2 have one stable attractor at high values while PU.1 has one stable attractor at low values. At the bifurcation point the GATA1 and GATA2 factors have one stable attractor at low values and one unstable at higher values while PU.1 has an unstable attractor at low values and a stable one at higher values.

In contrast, when varying the self-interaction strength for PU.1 we observe a non-linear behaviour. For low values of this strength, PU.1 exhibits one stable attractor with low values while GATA1 and GATA2 have one stable attractor at higher values. From the bifurcation point, PU.1 has two attractors: one unstable at low values and one stable at higher values of expression. At the bifurcation point for GATA1 and GATA2, the high expression attractor becomes unstable and a new stable attractor towards low values of expression appears (Figure 2B). Finally, we varied the self-interaction parameter of GATA2 and assessed the number and stability of the equilibrium points. For low values of self-interaction, GATA2 and GATA1 exhibit one stable attractor at low values while PU.1 shows one attractor at higher values. At the bifurcation point the low stable attractors of GATA1 and GATA2 become unstable and a new stable attractor emerges at higher values for both genes. For PU.1 the high value stable attractor becomes unstable and a new attractor at low values emerges (Figure 2C).

These findings identify GATA2 and PU.1 as potential important players in the process of erythroid lineage specification. Somewhat unexpectedly, GATA1 does not exhibit strong non-linear behaviour, suggesting it may not be of major relevance for the switching behavior required in lineage specification mechanisms.

### Knockout simulations suggest key role for GATA2 in lineage specification

By performing knockout (KO) simulations, we tested the functional impact of the absence of one regulator in the triad, in the normal time-evolution of the other two during erythroid differentiation (Figure 3). Although this is a simple circuit, given the relevance of each regulator for different cell states we consider their relative expression levels as indicative of lineage specification. GATA1 KO does not seem to have any significant impact on the normal expression of the other two regulators, suggesting that this regulator may not be relevant in the first instances of erythroid specification. This result is consistent with the bifurcation analysis, and surprising given previous reports relating the GATA1-PU1 axis to commitment decisions. GATA2 KO, on the other hand, has clear impact on the expression of the other two genes, particularly in the case of PU.1, which dramatically increases within the first few hours. GATA1 on the other hand, still maintains its broad qualitative behavior, but at lower relative levels. Finally, PU.1 KO leads to the sharp up-regulation of both GATA2 and GATA1 within the first few hours. Overall, these results suggest that the GATA2-PU.1 axis may be more relevant for committing cells to the erythroid lineage, specifically through the repression of PU.1 by GATA2. In this framework GATA2, and not GATA1, would act as the key regulator of lineage specification.

**Figure 3:**
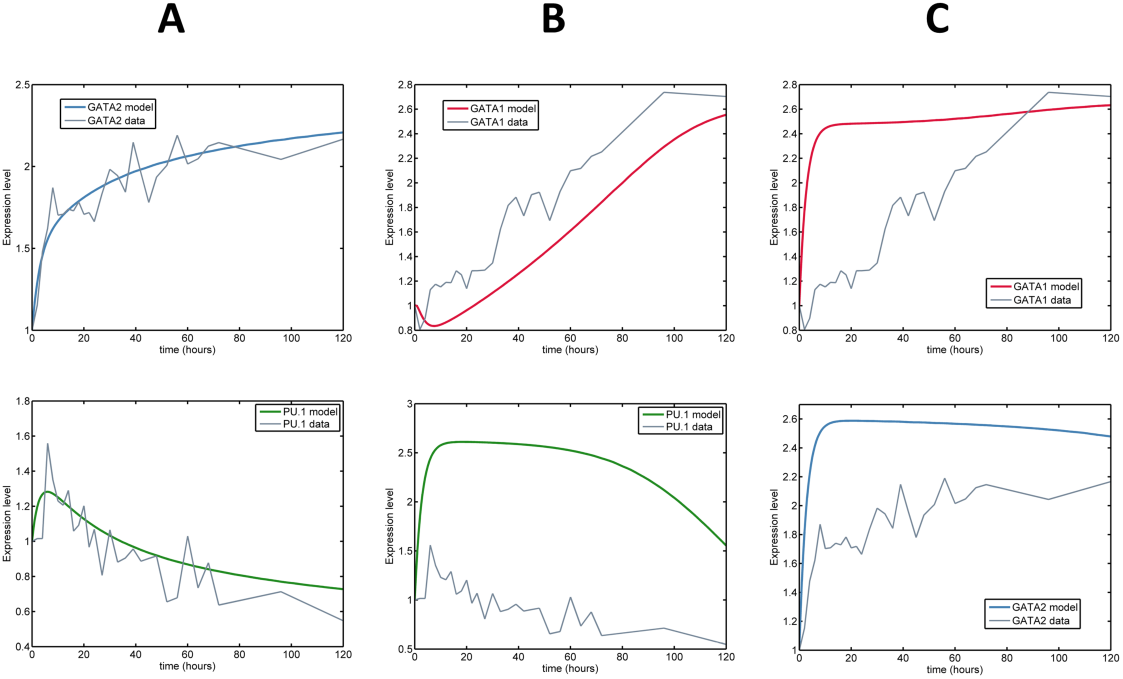
Expression time series of GATA1, PU.1 and GATA2 from modeling the triad architecture when simulating knockout of individual regulators. Experimental time series data are shown in grey. (A) GATA2 (blue) and PU.1 (green) when GATA1 is knocked out. No significant impact on GATA2 and PU.1 is observed. (B) GATA1 (red) and PU.1 when GATA2 is knocked out. A decrease of GATA1 is observed. Also, PU.1 increases substantially. (C) GATA1 and GATA2 when PU.1 is knocked out. An immediate increase of GATA1 and GATA2 is observed. Expression levels are normalized by expression at t=0 (multipotent state).

### Expansion of the GATA1/GATA2/PU.1 circuit provides insight into regulatory structure of early differentiation

In order to explore the mechanisms of early differentiation, and how commitment decisions effected within the GATA1/GATA2/PU.1 circuit propagate in the regulatory network, we extended the triad and included two downstream regulators, FOG1 and CEBPA, traditionally affiliated to the erythroid and myeloid lineages, respectively (Figure 4A). Reconstruction of this network was based on existing literature, as well as gene-expression and binding data provided by the same study that allowed inference of the triad sub-network. Data shows the existence of regulatory interactions between GATA2 and FOG1, as well as between PU.1 and FOG1 and PU.1 and CEBPA. Although ChIPSeq data provides directionality of these interactions, it cannot elucidate on their nature. We inferred the most likely set of interactions for this extended network by selecting the architectures that are best able to recapitulate experimentally observed gene expression for the five regulators (Figure 4B). According to our results, FOG1 is activated by GATA2 and repressed by PU1, which in turns activates CEBPA. These findings are consistent with current views of differentiation, where mutually antagonistic master regulators not only repress each other, but also bind to sets of downstream target genes, activating molecular programs of differentiation of their own lineage, while at the same time repressing programs of alternative lineages.

**Figure 4:**
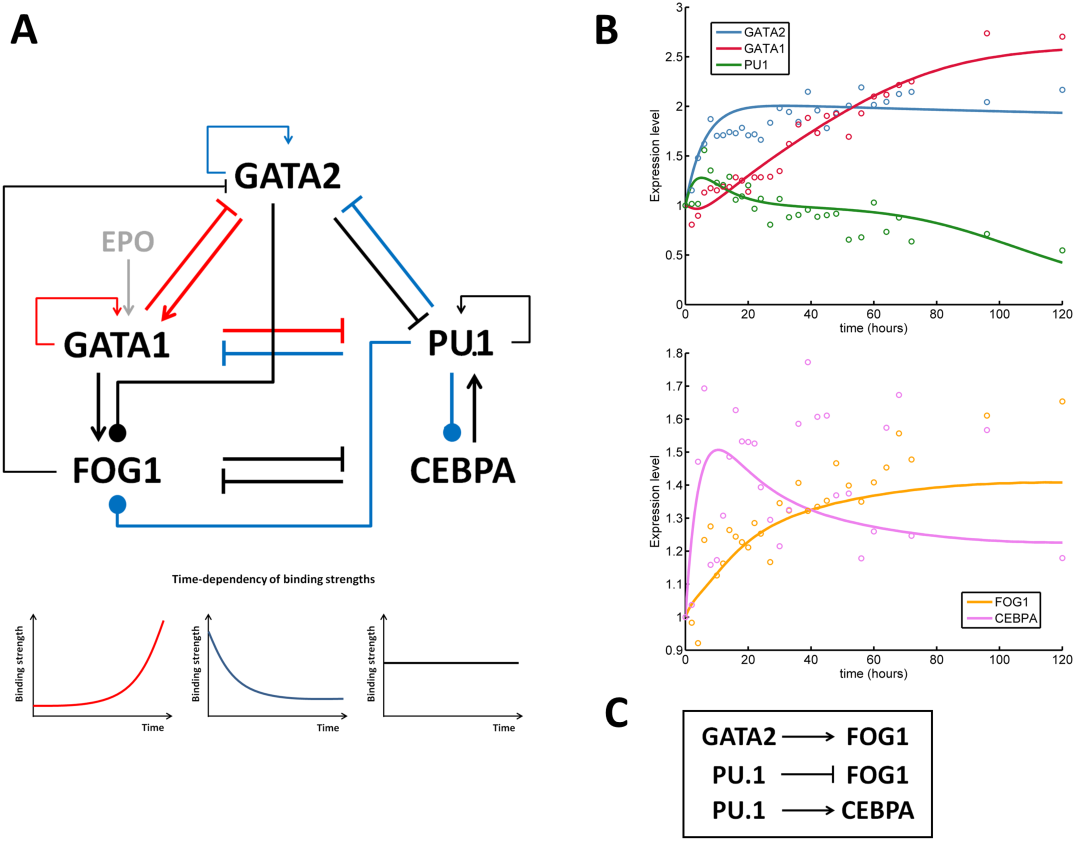
The extended interaction circuit and most likely interactions from fitting to experimental data. (A) The gene network for the mutual and self-regulatory interactions between GATA1, PU.1, GATA2, FOG1 and CEBPA. Red interaction indicates increase in time of the binding strength, blue corresponds to decreasing and black shows constant binding strengths. Interactions depicted by circle-end lines denote unknown type of interaction. (B) Time series of GATA1 (red), PU.1 (green), GATA2 (blue), FOG1 (orange) and CEBPA (magenta) from deterministic dynamics with the best set of estimated parameters. Solid lines represent simulation data and experimental data points are represented by circles. Expression levels are normalized by expression at t=0 (multipotent state). (C) The most likely type of regulatory interactions predicted for the unknown dynamics in the extended circuit.

### Knockout simulations in the extended network suggest combined regulation by GATA2 and GATA1 as an important mechanism of early differentiation

Using the same knockout simulation protocol as before, we analyzed how the absence of each regulator in the triad impacts on the process of early erythroid differentiation (Figure 5). Although this process is necessarily much more complex, involving a large number of genes, we consider the regulation of FOG1 and CEBPA as an illustration of how changes in master regulators can be carried out in lower levels of the network.

**Figure 5:**
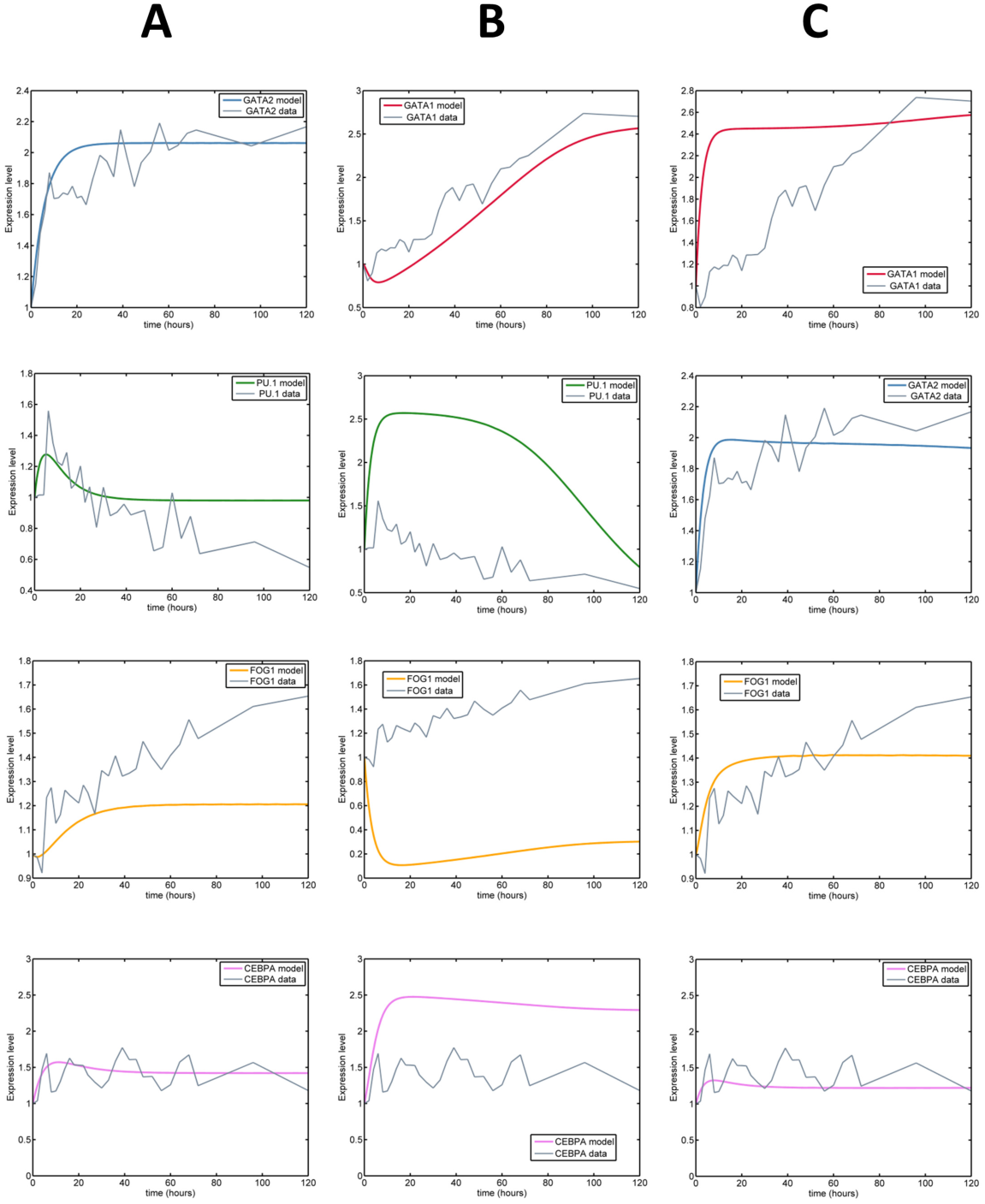
Expression time series of GATA1, PU.1, GATA2, FOG1 and CEBPA from modeling the extended architecture in Figure 4A when simulating GATA1, GATA2 and PU.1 knockouts. Experimental data are shown in grey. (A) GATA2 (blue), PU.1 (green) FOG1 (orange) and CEBPA (magenta) when GATA1 is knocked out. The impact of the GATA1 knockout is a decrease of FOG1 and a slight increase of PU.1. (B) GATA1 (red), PU.1, FOG1 and CEBPA when GATA2 is knocked out. A decrease of GATA1 and FOG1 is observed. Also, PU.1 and CEBPA increase substantially. (C) GATA1, GATA2, FOG1 and CEBPA when PU.1 is knocked out. An immediate increase of GATA1, GATA2 and FOG1 and a slight decrease in CEBPA are observed. Expression levels are normalized by expression at t=0 (multipotent state).

GATA1 KO exhibits the same weak influence over the expression of GATA2 and PU.1 as before. However, it now affects its downstream target FOG1, which reaches a plateau at levels lower than expected in a normal differentiation time-course. Despite this perturbation, CEBPA expression remains stable.

GATA2 KO results are also consistent with the previous observations within the triad, with both GATA1 and more so PU.1 expression being affected. Interestingly, this perturbation propagates to the lower level of regulation, causing the down-regulation of FOG1 and the up-regulation of CEBPA. The fact that GATA2 is directly connected to FOG1 and has a strong impact on PU.1, which in turn regulates CEBPA, explains this strong effect, suggesting potential significant effects in terms of lineage differentiation.

Finally, PU.1 KO also affects expression of GATA1 and GATA2 in approximately the same measure as before, but seems not to have significant impact in the next level of regulation with minor changes observed in the expressions of FOG1 and CEBPA.

Overall, these results show that within the triad of master regulators, GATA2 has the strongest influence over the extended network, with absence of expression causing a perturbation that propagates to the lower level of regulation. One of the suggested mechanisms for driving erythroid differentiation is the process of GATA switching, in which GATA2 initially binds to and regulates a large number of genes, and is gradually replaced by GATA1 as driver of the process. Our recent experimental data suggests that in fact binding of both GATA2 and GATA1 may be necessary in many instances, with GATA2 acting as a facilitator of GATA1 binding, but instead of being replaced, acting together during differentiation. The regulation of FOG1 in our system provides a simple example to address this question. We observe that in the absence of GATA1, FOG1 experiences early up-regulation but cannot reach normal levels throughout the time-course. On the other hand, in the absence of GATA2, FOG1 is down-regulated altogether, falling below the observed expression levels in the multipotent state. These results are consistent with the view that both GATA2 and GATA1 are necessary for normal regulation of some downstream targets.

## Discussion

Inspired by recent high-resolution gene expression time-course and ChIPSeq binding data, we addressed fundamental aspects of the regulatory mechanisms underlying erythroid lineage specification and differentiation, through the use of dynamical modeling. In particular, we focused on the triad of genes composed by GATA1/GATA2/PU1 and investigated its potential for switch-like behavior, and the impact of decisions effected within this core circuit in the regulation of downstream transcriptional programs.

Bistability is a key property of regulatory circuits in differentiating systems, providing memory and robustness to external fluctuations, and allowing the generation of phenotypic diversity [14]. The relevance of bistable switches has been modeled and demonstrated for a number of differentiation systems ([15], and references therein). In the erythroid lineage, previous studies have theoretically deduced that bistability can emerge along the GATA1/PU.1 axis [7, 9]. A subsequent modeling effort has suggested that, considering a detailed description of the interactions between these two genes, the GATA1/PU.1 pair is not sufficient to generate bistability by itself, and a third element must be involved [8]. In this work, we revisited the paradigmatic GATA1/PU.1 switch in order to characterize its bistability potential, in view of our recent findings in [4], suggesting the importance of PU.1 regulation by GATA2 in erythroid lineage specification. By varying the self-interactions one at a time for GATA1, GATA2 and PU.1, we concluded that the self-interactions of GATA2 and PU.1 lead to one unstable and one stable fixed point. In contrast, this was not the case for GATA1, where a very smooth behavior was observed. This suggests that, when constraining system parameters to experimental data, bistability cannot be observed within the GATA1-PU.1 axis, and instead GATA2 seems to have the potential to regulate cell fate decisions. This picture could also be observed when including the downstream regulators FOG1 and CEBPA (results not shown).

Using the same high-resolution experimental data and inference strategy as in [4], we expanded the initial GATA1/GATA2/PU.1 sub-network to include the downstream regulators FOG1 and CEBPA, affiliated with the erythroid and myeloid lineages respectively. Differentiation into different lineages has been reported to proceed by sequential decision points, where cells engage in a series of binary fate decisions governed by pairs of master regulators [5]. The regulatory actions of these transcription factors is not confined to each other, since they can bind to large sets of downstream genes, coordinating entire transcriptional programs that drive differentiation. For our network, ChIPSeq data could provide the directionality but not the nature of interactions between PU.1 and FOG1; PU.1 and CEBPA; and GATA2 and FOG1. Since these configure regulatory interactions between master regulators and downstream lineage-affiliated genes, they provide a simple testing ground for the existence of regulatory program coordination by regulators of the core network. We generated the 8 possible architectures involving these 3 unknown interactions and observed that the best fits to gene expression time series data were provided by the configuration where PU.1 represses FOG1 and activates CEBPA, whereas GATA2 activates FOG1. These observations are in line with current views that regulators of lineage specification act on downstream levels of the newtork, by activating genes affiliated with their own lineage and repressing those affiliated with alternative fates. A recent study has experimentally observed this pattern in erythroid differentiation, for a core network composed by GATA1, SCL and KLF1 [11]. The authors also suggest, by means of a mathematical modeling, that this dual mechanism of repression of both the antagonistic master regulator and its affiliated genes provides a robust mechanism for lineage specification. Our results also confirm the applicability of the network inference approach developed in [4], combining dynamical modeling and experimental data to build upon and expand known regulatory circuits.

Based on both the triad and the extended network circuits, we performed simulations of knockout experiments in order to test the functional relevance of individual regulators in the processes of lineage specification and early differentiation. Concerning early events confined to the triad, we could confirm our earlier observations in the bifurcation studies. In particular, we observed that GATA1 knockout did not have signifi-cant impact on the expression of the other two regulators, particularly when compared to PU.1 knockout and more so GATA2 knockout. Overall, these results suggest a key role for GATA2 in the regulatory events of lineage specification, in particular through the GATA2-PU.1 axis. Results were consistent when performing the simulations in the triad-only or extended models.

When expanding our observations to the downstream level of regulation, we observed that GATA2 knockout still had the strongest impact, with the levels of FOG1 and CEBPA clearly suggesting propagation of upstream perturbation events. PU.1 knockout on the other hand, did not seem to propagate to either CEBPA or FOG1. Interestingly, GATA1 knockout clearly affects downstream expression of FOG1, suggesting that despite its weak impact on the other two triad regulators, GATA1 is relevant for subsequent regulatory events, in accordance with its recognized key role in erythroid differentiations. We dissected the combined regulation of FOG1 by both GATA1 and GATA2 in order to gain some basic insight into potential mechanisms underlying GATA switching. Our results suggest that both GATA2 and GATA1 are needed in order to recapitulate the observed gene expression time-course for FOG1. This is consistent with the recent experimental observations in [4], which suggest that binding of both GATA2 and GATA1 may be necessary throughout the entire erythroid differentiation, with GATA2 acting as a facilitator of GATA1 binding. This is in contrast with more traditional views of GATA switching, where GATA2 is assumed to be replaced, and transfer control to GATA1 [12, 13].

## Methods

### Models framework

To elucidate the role played by each regulator in the GATA1/GATA2/PU.1 sub-network, we expanded the sub-network inferred in [4], by including the downstream regulators FOG1 and CEBPA. As in [4], we use a mechanistic model of deterministic rate equations that bridges ChIPseq binding data with gene expression time series data to determine the nature of the interactions within this extended sub-network.

Our results from [4] along with existing literature lead to the following assumptions:

- GATA1 and PU.1 exhibit the well-established cross-inhibition and self-activation structure [5, 8, 10]. Additionally, we assume heterodimer formation involving GATA1 and PU.1 whenever binding occurs [16].
- GATA2 self-activates, establishes a negative feed-back circuit with GATA1, and cross-inhibition with PU.1 [4].
- CEBPA self-activates [17], exhibits cross-inhibition with FOG1 [18, 8] and activates PU.1 [6].
- FOG1 is activated by GATA1 [8, 6] and represses GATA2 [19].
- From our ChIPSeq data, GATA2 interacts with FOG1, and PU.1 interacts with both CEBPA and FOG1 but the exact nature of these interactions is unknown.
- In order to capture the culture conditions in erythroid differentiation, Epo is present as an external factor, impinging on GATA1 with constant activation strength.
- As in [4], we defined the binding strength behaviors in time as constant or exponentially increasing/decreasing, based on the qualitative behavior observed in ChIPSeq data, by comparing binding in the multipotential and erythroid-differentiated states.

## Mathematical formulation

### The GATA1/GATA2/PU.1 triad

The dynamics of the triad architecture in (Figure 1A) are given by a set of differential equations following the Shea-Ackers thermodynamic approach for transcription modeling [20, 21]. These equations have 18 unknown parameters, binding strengths and decay rates, which were estimated from gene expression time series data.

The equations describing the behavior of GATA1, PU.1 and GATA2 are then given by:

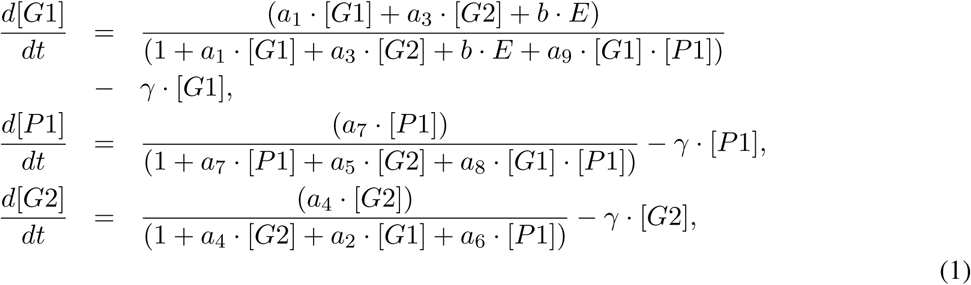

where the GATA1, GATA2, PU.1 and Epo concentration levels are denoted by [G1], [G2], [P1] and *E* respectively and *t* denotes the time. The varying binding strengths parameters are parameterized as

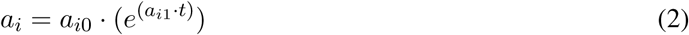

with the exception of *a*_5_ and *a*_7_ which are constant. The degradation rates are denoted by *γ* and we consider them to be similar for all three molecules and *b* represents the strength of Epo signaling on the system.

### Extended network inference and modeling

The extended network model consists of the GATA1/GATA2/PU.1, inheriting the molecular interactions described above, and including FOG1 and CEBPA. The added downstream regulators were linked to the existing network according to existing literature and our ChIPSeq data, as described above. However, from the ChIPSeq data one cannot infer the nature of interactions between GATA2 and FOG1, as well as PU.1 with FOG1 and CEBPA. Hence we have explored all possible sign combination for the three interactions, in order to infer the most likely regulatory effect, activation or repression. Through dynamical modeling, we then assessed which of the 8 resulting possible architectures is the most accurate in explaining the observed gene expression time series behavior for the five genes. Again, the Shea-Ackers formalism [20, 21] is used for the dynamics. Each set of resulting differential equations has 29 parameters corresponding to binding strengths and decay rates. The binding strength variations observed from the data were again modeled as exponential increase or decay in time of the corresponding parameter value. If the data showed no significant variations of the binding strength then the corresponding parameter value was considered constant.

The extended architecture that was able to best describe the five gene expression time series data was modeled using the following set of equations:

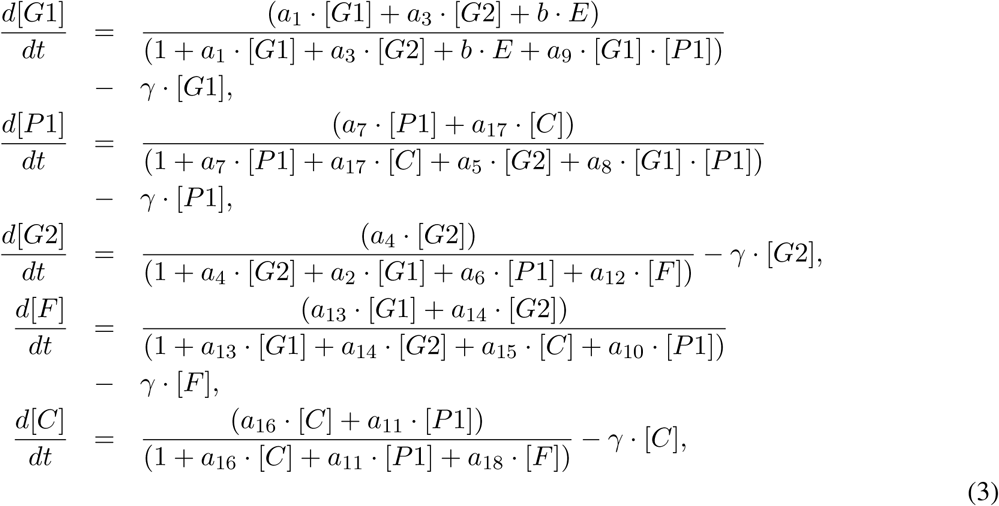

where the GATA1, PU.1, GATA2, FOG1, CEBPA and Epo concentration levels are denoted by [G1], [P1], [G2], [F], [C] and *E* respectively and *t* denotes the time. Again, varying binding strength parameters were modeled as exponential functions as described in (Equation 2) except *a*_5_, *a*_7_ and *a*_12*…*18_ which are constant. The degradation rate is denoted by *γ* and considered to be similar for all five molecules. As above, *b* represents the strength of Epo signaling on the system.

The simulated annealing algorithm [22] was used iteratively to estimate the best parameter set of each of the 8 proposed dynamical models. The aim is to find the parameter set *θ*_*i*_, *i* = 1*…*8 that minimizes the quadratic error *E*_*i*_ between model predictions and the microarray time-series data consisting of 27 timepoints covering the time period from 0 to 120 hours of erythroid differentiation.

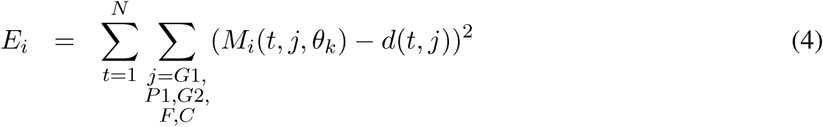

where *M*_*i*_(*t, j, θ*_*k*_) denotes model generated points, *d*(*t, j*) represent the experimental data points normalized by expression value at time *t* = 0, *i* is the architecture, *t* is the time and *j* denotes the five different genes, *k* = 1 : 29 represents the parameter index. The global minimum of *E*_*i*_ was approximated using the simulated annealing algorithm. The parameter estimation process is an iterative process where we run the optimization protocol for multiple times keeping the set of parameters that minimizes *E*_*i*_, each time. The results are then used for calculating the scores *S*_*i*_ for each architecture. The scores are used to identify the best architecture describing the microarray gene expression time series data. Architecture scores represent the ratio between the number of estimation runs *r*_*i*_ that produced solutions with values of *E*_*i*_ under a specified threshold *τ* and the total number of runs for each architecture *R*_*i*_.

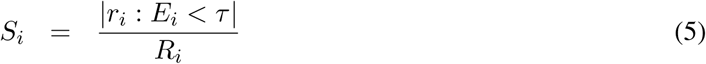

### Bifurcation study

The steady state solutions are analyzed as a function of the network parameters, in particular, the self-interaction binding strengths of GATA1, PU.1 and GATA2. Local bifurcation analyses were carried out by assessing the changes in the stability and the number of the equilibrium points when the values of the parameters corresponding to the self-interaction binding strengths of the genes from the upstream triad were varied while the rest of parameters were kept fixed at their values at time *t* = 0. The bifurcation diagrams were generated using Oscill8 (http://sourceforge.net/projects/oscill8).

### 0.1 Knock out simulation

We perform simple approximations for gene knockout experiments by fixing the knocked out gene expression value to 0 throughout the simulation. Gene expression for each regulator was initialized with a value of 0 and the equation describing its evolution was replaced by the constant 0 throughout. These simulations cannot be directly equated to traditional experimental knockout experiments, and are more akin to knockdown methods, where gene expression is targeted and reduced by means of techniques such as lentiviral-mediated RNA interference. In such a framework, our simulations configure the hypothetical scenario of 100% efficiency, leading to complete absence of expression of the targeted gene. These simulations provide a simple tool for exploring *in silico* loss-of-function scenarios in the context of erythroid lineage specification and early differentiation.

Inference of parameters and simulations of the differential equations were implemented using MATLAB software (The Mathworks).

## 1 Acknowledgments

The authors thank Gill May, Alex Tipping and Vijay Chickarmane for valuable discussions. JT is part of the PhD Program in Computational Biology at Instituto Gulbenkian de Ciencia, Oeiras, Portugal, funded by Fundacao para a Ciencia e Tecnologia (SFRH/BD/33208/2007). This work was funded by the Swedish Foundation for Strategic Research, Senior Individual Grant (CP) and the Swedish Research Council (VO, JT and CP)

